# FlyBrainLab: Accelerating the Discovery of the Functional Logic of the *Drosophila* Brain in the Connectomic/Synaptomic Era

**DOI:** 10.1101/2020.06.23.168161

**Authors:** Aurel A. Lazar, Tingkai Liu, Mehmet Kerem Turkcan, Yiyin Zhou

## Abstract

In recent years, a wealth of *Drosophila* neuroscience data have become available. These include cell type, connectome and synaptome datasets for both the larva and adult fly. To facilitate integration across data modalities and to accelerate the understanding of the functional logic of the fly brain, we developed an interactive computing environment called FlyBrainLab.

FlyBrainLab is uniquely positioned to accelerate the discovery of the functional logic of the *Drosophila* brain. Its interactive *open source* architecture seamlessly integrates and brings together computational models with neuroanatomical, neurogenetic and electrophysiological data, changing the organization of neuroscientific fly brain data from a group of seemingly disparate databases, arrays and tables, to a well structured data and executable circuit repository.

The FlyBrainLab User Interface supports a highly intuitive and automated workflow that streamlines the 3D exploration and visualization of fly brain circuits, and the interactive exploration of the functional logic of executable circuits created directly from the explored and visualized fly brain data. Furthermore, efficient comparisons of circuit models are supported, across models developed by different researchers, across different developmental stages of the fruit fly and across different datasets.

The FlyBrainLab Utility Libraries help untangle the graph structure of neural circuits from raw connectome and synaptome data. The Circuit Libraries facilitate the exploration of neural circuits of the neuropils of the central complex and, the development and implementation of models of the adult and larva fruit fly early olfactory systems.

Seeking to transcend the limitations of the connectome, FlyBrainLab provides additional libraries for molecular transduction arising in sensory coding in vision and olfaction. Together with sensory neuron activity data, these libraries serve as entry points for discovering circuit function in the sensory systems of the fruit fly brain. They also enable the biological validation of developed executable circuits within the same platform.

## Introduction

The era of connectomics/synaptomics ushered in the advent of large-scale availability of highly complex fruit fly brain data [1], [2], [3], [4], while simultaneously highlighting the dearth of computational tools with the speed and scale that can be effectively deployed to uncover the functional logic of fly brain circuits. In the early 2000’s, automation tools introduced in computational genomics significantly accelerated the pace of gene discovery from the large amounts of genomic data. Likewise, there is a need to develop tightly integrated computing tools that automate the process of 3D exploration and visualization of fruit fly brain data with the interactive exploration of executable circuits.

To meet this challenge we have built an open source interactive computing platform called FlyBrainLab. FlyBrainLab is uniquely positioned to accelerate the discovery of the functional logic of the Drosophila brain. It is designed with three main capabilities in mind: 1) 3D exploration and visualization of fruit fly brain data, 2) creation of executable circuits directly from the explored and visualized fly brain data in step 1), and 3) interactive exploration of the functional logic of the executable circuits devised in step 2) (see Figure 1).

**Figure 1:**
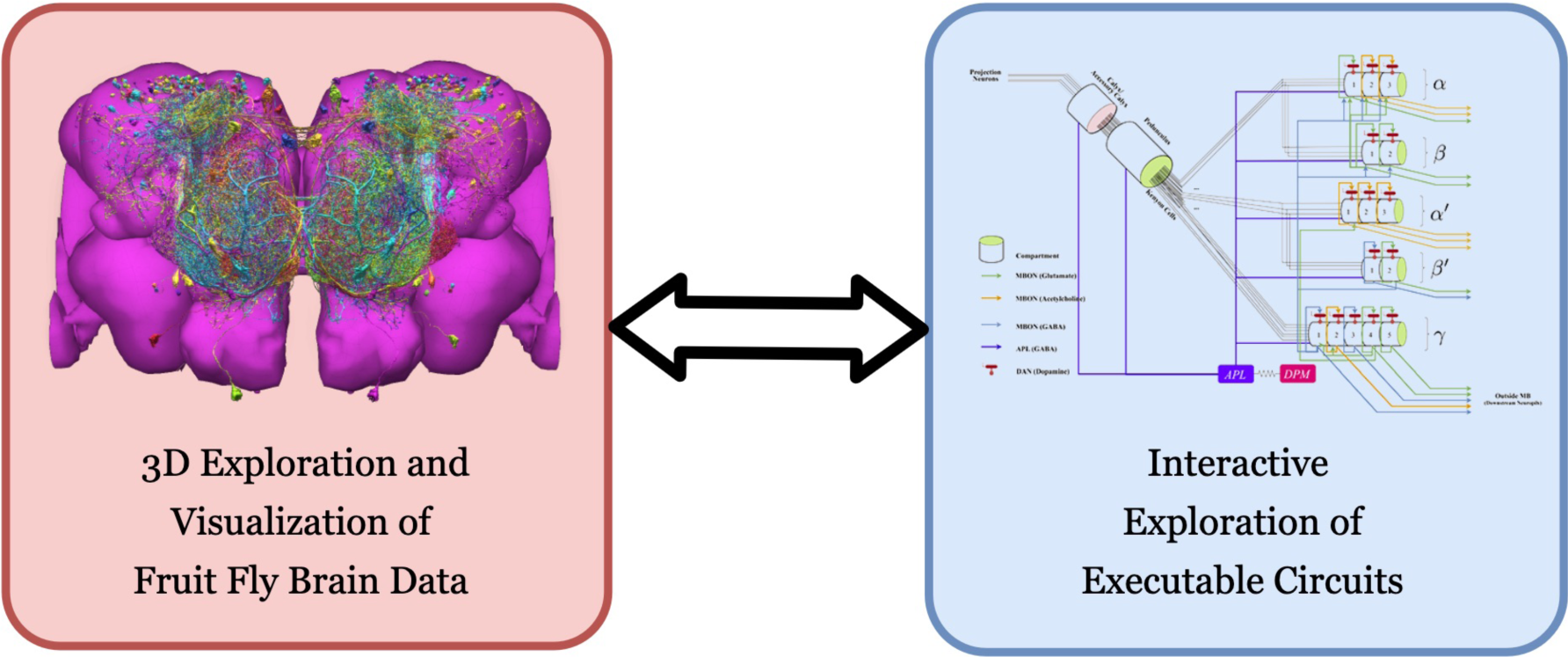
FlyBrainLab provides, within a single working environment, 3D visualization and exploration of connectome data, interactive exploration of executable circuit diagrams and real-time code execution.

To achieve a tight integration of the three main capabilities into the single working environment sketched in Figure 1, FlyBrainLab exhibits an architecture (see Supplementary Figure **1**) with a number of key backend services provided by the NeuroArch Database [5] and the Neurokernel Execution Engine [6], and the NeuroMynerva front-end supporting an integrated 3D graphics user interface (GUI) (see also Supplementary Figure **2**). The key feature of the FlyBrainLab architecture is the tight integration between the NeuroArch and Neurokernel components that is further reinforced by NeuroMynerva. The tight integration is critical for achieving the automated exploration of fly brain data and the rapid creation and exploration of the function of executable circuits.

To accelerate the generation of executable circuits from biological data, NeuroMynerva supports the following workflow (see also Supplementary Figure **2**).

First, the 3D GUI (see Supplementary Figure **2**, top left) supports the visual exploration of fly brain data, including neuron morphology, synaptome, and connectome from all available data sources, stored in the custom-built NeuroArch Database [5]. With plain English queries (see Supplementary Figure **2**, top left), a layperson can perform sophisticated database queries with only knowledge of fly brain neuroanatomy [7].

Second, the circuit diagram GUI (see Supplementary Figure **2**, top right) enables the interactive exploration of executable circuits stored in the NeuroArch Database. By retrieving tightly integrated biological data and executable circuit models from the NeuroArch Database, NeuroMynerva supports the interaction and interoperability between the biological circuit built for morphological visualization and the executable circuit created and represented as an interactive circuit diagram, and allows them to build on each other. This helps circuit developers to more readily identify the modeling assumptions and the relationship between neuroanatomy, neurocircuitry and neurocomputation.

Third, the Neurokernel [6] Execution Engine provides circuit execution on multiple computing nodes/GPUs. The tight integration in the database also allows the execution engine to fetch executable circuit directly from the NeuroArch Database. The tight integration between NeuroArch and Neurokernel is reinforced and made user transparent by NeuroMynerva.

Finally, the GUIs can operate in tandem with command execution in Jupyter notebooks (see also Supplementary Figure **2**, bottom center). Consequently, fly brain circuits and circuit diagrams can be equivalently processed using API calls from Python, thereby ensuring the reproducibility of the exploration of similar datasets with minimal modifications.

Analysis, evaluation and comparison of circuit models, either among versions developed by one’s own, or among those published in literature, are often critical steps towards discovering the functional logic of brain circuits. Three types of analyses, evaluations and comparisons are of particular interest. First, starting from a given dataset and after building a number of circuit models published in the literature, analyze and compare them under the same evaluation criteria. Second, automate the construction of executable circuits from datasets gathered by different labs and, analyze, evaluate and compare the different circuit realizations. Third, analyze, evaluate and compare fruit fly brain circuits at different developmental stages. In what follows, we present results, supported by the FlyBrainLab Circuits Libraries (see Methods), demonstrating the comprehensive evaluation, analysis and comparison capability of FlyBrainLab.

## Results

### Analyzing, Evaluating and Comparing Circuit Models of the Fruit Fly Central Complex

We first demonstrate the workflow supported by FlyBrainLab for analyzing, evaluating and comparing circuit models of the fruit fly Central Complex (CX) based on the FlyCircuit dataset [1]. The circuit connecting the ellipsoid body (EB) and the protocerebral bridge (PB) in the CX has been shown to exhibit ring attractor dynamics [11–13]. Recently, a number of researches investigated circuit mechanisms underlying these dynamics. Here, we developed a CXcircuits Library for analyzing, evaluating and comparing various CX circuit realizations. Specifically, we implemented 3 of the circuit models published in the literature, called here model A [8], model B [9], and model C [10], and compared them in the same FlyBrainLab programming environment.

In Figure 2(a1, b1, c1), the anatomy of the neuronal circuits considered in model A, B and C, is respectively depicted. The corresponding interactive circuit diagram is shown in Figure 2(a2, b2, c2). Here, model A provides the most complete interactive CX circuit, including the core subcircuits for characterizing the PB-EB interaction with the EB-LAL-PB, PB-EB-LAL, PB-EB-NO, PB local and EB ring neurons (see Methods and [8] for commonly used synonyms). Models B and C exhibit different subsets of the core PB-EB interaction circuit in model A. While no ring neurons are modeled in model B, PB local neurons are omitted in model C. They, however, do no model other neurons in the CX, *e*.*g*., those innervate the FB.

**Figure 2:**
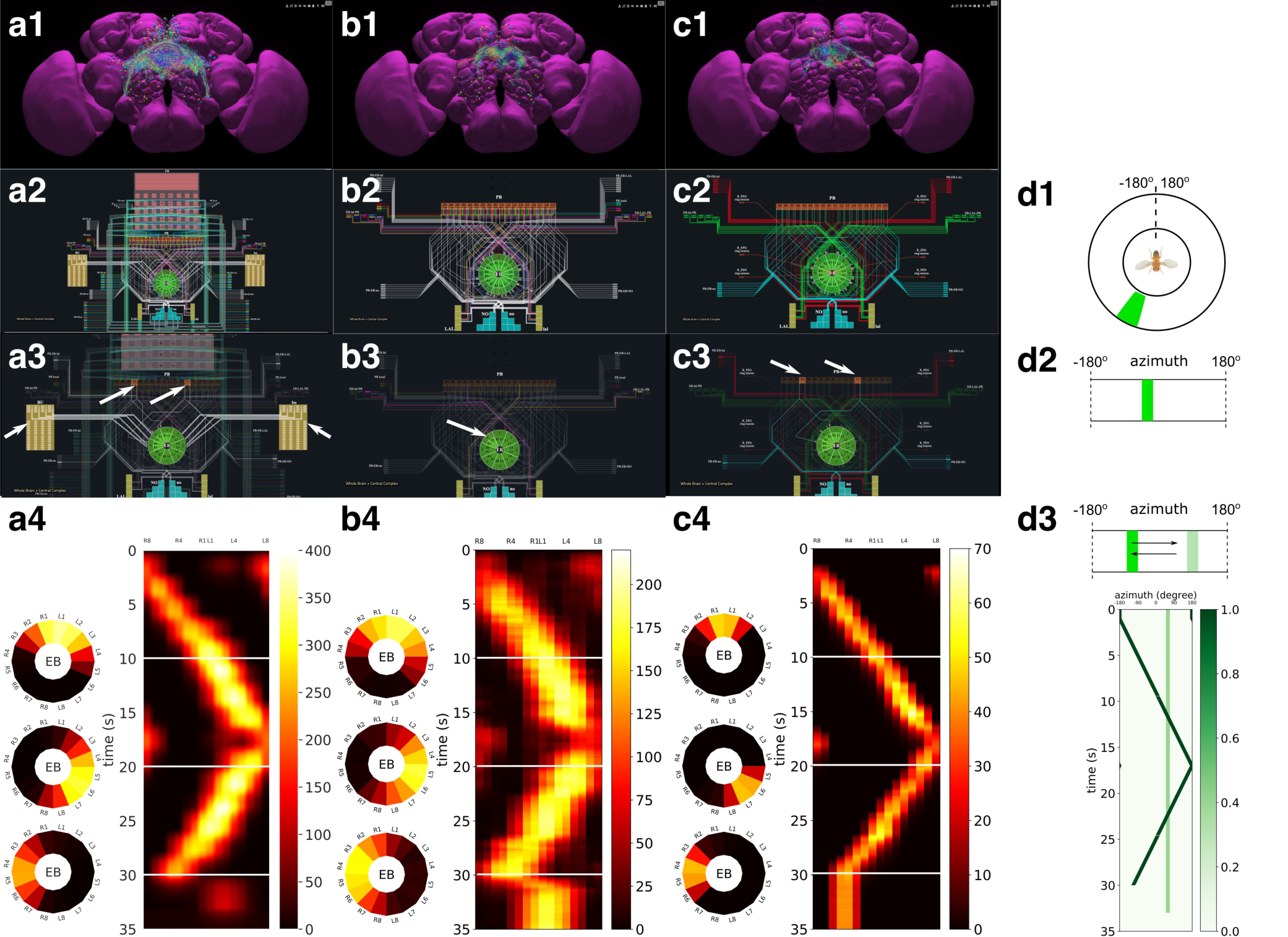
Analysis, evaluation and comparison of 3 models of CX published in the literature. (a) Model A [8], (b) Model B [9], (c) Model C [10]. (a1, b1, c1) Morphology of the neurons visualized in the NeuroNLP window (see Supplementary Figure **2**). (a2, b2, c2) Neuronal circuits in the NeuroNLP window depicted in the NeuroGFX window (see Supplementary Figure **2**) as abstract interactive circuit diagrams. The naming of the ring neurons in (c2) follows [10]. (a3, b3, c3) When a single vertical bar is presented in the visual field (d1/d2), different sets of neurons/subregions (highlighted) in each of the models, respectively, receive either current injections or external spike inputs. (a4, b4, c4) The mean firing rates of the EB-LAL-PB neurons innervating each of the EB wedges of the 3 models (see Methods), in response to the stimulus shown in (d3). Insets show the rates at 10, 20, and 30 seconds, respectively, overlaid onto the EB ring. (d1) A schematic of the visual field surrounding the fly. (d2) The visual field flattened. (d3) Input stimulus consisting of a bar moving back and forth across the screen, and a second fixed bar at 60*°* and with lower brightness.

In Supplementary Video 1, we demonstrate the interactive capabilities of the three models side-by-side, including the visualization of the morphology of CX neurons and the corresponding executable circuits, user interaction with the circuit diagram revealing connectivity pattern, and the execution of the circuit. In the video, the visual stimulus depicted in Figure 2(d3) was presented to all three models (see Methods for the details of generating the input stimulus for each model). The responses, measured as the mean firing rate of EB-LAL-PB neurons within contiguous EB wedges, are shown in Figure 2(a4, b4, c4), respectively. Insets depict the responses at 10, 20 and 30 seconds. During the first second, a moving bar in its fixed initial position and a static bar are presented. The moving bar displays a higher brightness than the static bar. All three models exhibited a single-bump (slightly-delayed) response tracking the position of the moving bar. The widths of the bumps were different, however. After 30 seconds, the moving bar disappeared and models A and B shifted to track the location of the static bar, whereas the bump in model C persisted in the same position where the moving bar disappeared. Furthermore, for models B and C but not for model A, the bumps persisted after the removal of the visual stimulus (after 33 seconds), as previously observed *in vivo* [11, 12].

By comparing these circuit models, we notice that, to achieve the ring attractor dynamics, it is critical to include global inhibitory neurons, *e*.*g*., PB local neurons in models A and B, and ring neurons in models A and C. The model A ring neurons featuring a different receptive field and the ring neurons in model C receiving spike train input play a similar functional role. However, to achieve the ring attractor dynamics characterized by a single bump response to multiple bars and persistent bump activity after the removal of the vertical bar, model C only required three out of the five core neuron types (see Methods), whereas model B requires all all four neuron types included.

### Analyzing, Evaluating and Comparing of Adult Antenna and Antennal Lobe Circuit Models Based upon the FlyCircuit and Hemibrain Datasets

In the second example we demonstrate the effect on modeling the antenna and antennal lobe circuits due to, respectively, the FlyCircuit [1] and the Hemibrain [4] datasets (see also Methods).

We start by exploring and analyzing the morphology and connectome of the olfactory sensory neurons (OSNs), antennal lobe projection neurons (PNs) and local neurons (LNs) of the FlyCircuit [1] and the Hemibrain [4] datasets (see Figure 3). The high resolution electron microscopy reveals new connectivity motifs in the Hemibrain dataset between olfactory sensory neurons, projection neurons and local neurons (see Fig. 3(a)). Following [14], we first constructed the two layer circuit shown in Figure 3(b) (left) and then constructed a more extensive connectome/synaptome model of the adult antennal lobe shown in Fig. 3(b) (right).

**Figure 3:**
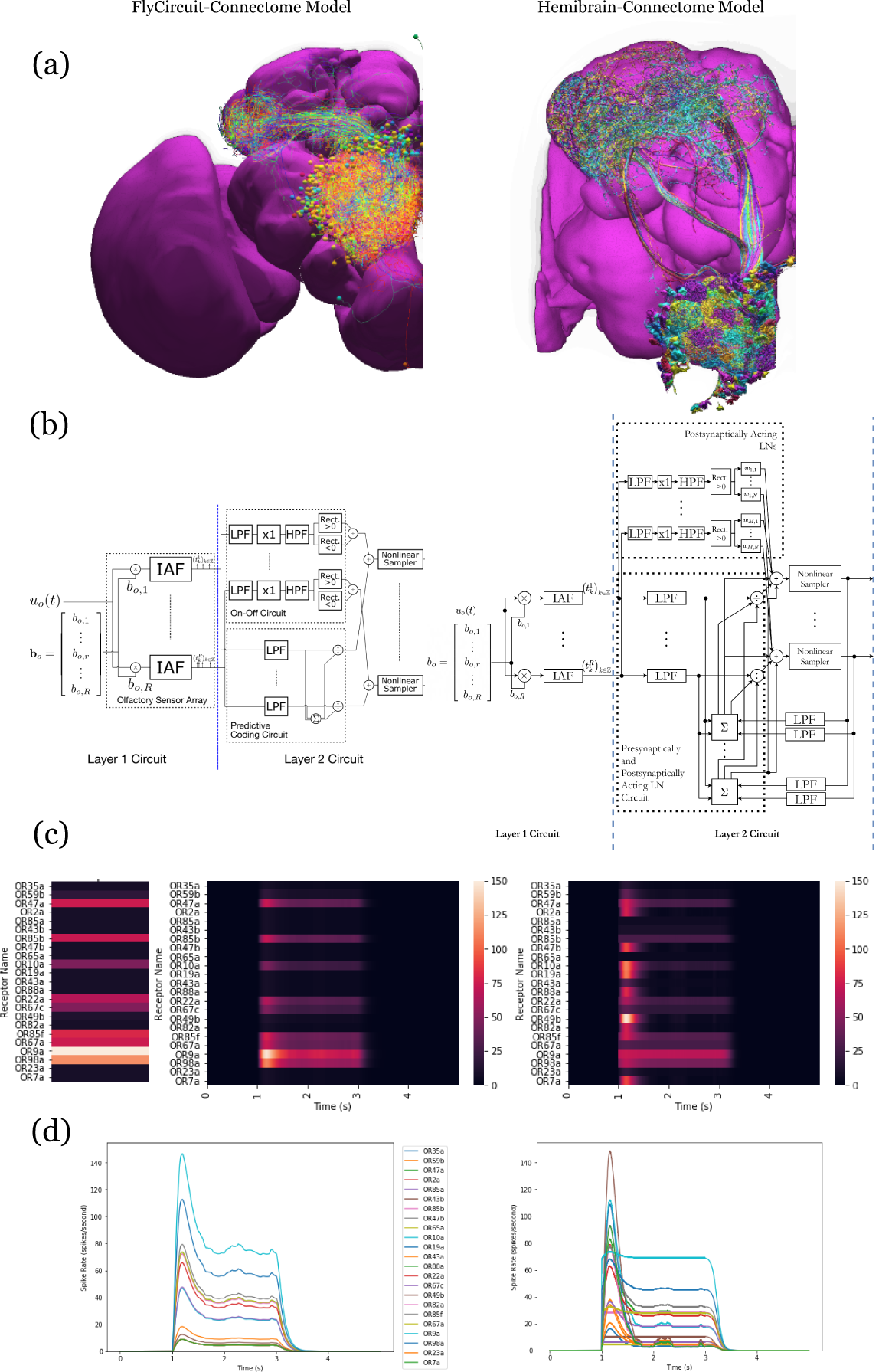
Analysis, evaluation and comparison between 2 models of the antenna and antennal lobe circuit of the adult fly based on the FlyCircuit (left) dataset [14] and an exploratory model based on the Hemibrain (right) dataset [4]. (a) Morphology of olfactory sensory neurons, local neurons and projection neurons in the antennal lobe for the two datasets. The axons of the projection neurons and their projections to the mushroom body and lateral horn are also visible. (b) Circuit diagrams depicting the antenna and antennal lobe circuit motifs derived from the two datasets. (c) Response of the antenna/antennal lobe circuit to a constant ammonium hydroxide step input applied between 1s and 3s of a 5 second simulation; (left) the interaction between the odorant and 23 olfactory receptors is captured as the vector of affinity values; (middle and right) a heatmap of the uniglomerular PN PSTH values (spikes/second) grouped by glomerulus for the 2 circuit models. (d) The PN response transients of the 2 circuit models for uniform noise input with a minimum of 0ppm and a maximum of 100ppm preprocessed with a 30Hz low-pass filter [15] and delivered between 1s and 3s.

Execution of and comparison of the results of these two circuit models show quantitatively different PN output activity in steady-state (Fig. 3(c)) and for transients (Fig. 3(d)). A prediction [14, 16] made by the antenna and antennal lobe circuit shown in Figure 3(b) (left) using the FlyCircuit data has been that the PN activity, bundled according to the source glomerulus, is proportional to the vector characterizing the affinity of the odorant-receptor pairs (Fig. 3(c), left column).

The transient and the steady state activity response are further highlighted in Figure 3(d) for different amplitudes of the odorant stimulus waveforms. The initial results show that the circuit on the right detects with added emphasis the beginning and the end of the odorant waveforms.

The complex connectivity between OSNs, LNs and PNs revealed by the Hemibrain dataset suggests that the adult antennal lobe circuit encodes additional odorant representation features [4].

### Analyzing, Evaluating and Comparing Early Olfactory Circuit Models of the Larva and the Adult Fruit Flies

In the third example, we investigate the difference in odorant encoding and processing in the *Drosophila* Early Olfactory System (EOS) at two different developmental stages, the adult and larva (see also Methods).

We start by exploring and analyzing the morphology and connectome for the Olfactory Sensory Neurons (OSNs), Antennal Lobe Projection Neurons (PNs) and Local Neurons (LNs) of the adult Hemibrain [4] dataset and the LarvaEM dataset (see Figure 4 (a)).

**Figure 4:**
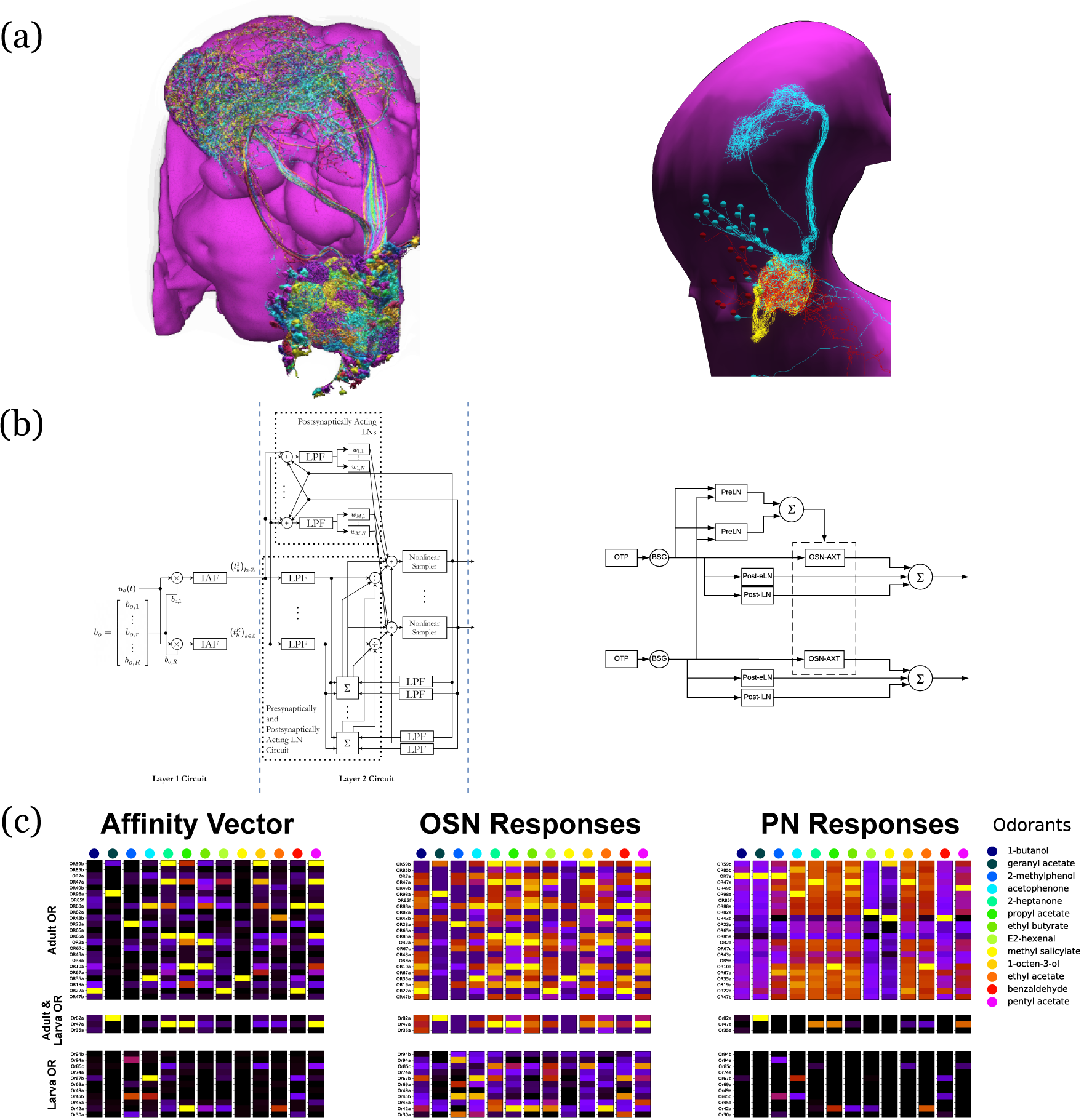
Evaluation and Comparison of two *Drosophila* Early Olfactory System (EOS) models describing adult (*left*, developed based on Hemibrain dataset) and larval (*right*, developed based on LarvaEM dataset) circuits. (a) Morphology of Olfactory Sensory Neurons (OSNs) in the Antenna (ANT), Local Neurons (LNs) in the Antennal Lobe (AL) and Projection Neurons in the AL. (b) Circuit diagrams depicting the Antenna and Antennal Lobe circuit motifs. (c) (left) Interaction between 13 odorants and 37 odorant receptors (ORs) characterized by affinity values. The ORs expressed only in the adult fruit flies are grouped in the top panel; the ones that are expressed in both the adult and the larva are grouped in the middle panel; and those expressed only in the larva are shown in the bottom panel. Steady-state outputs of the EOS models to a step concentration waveform of 100 ppm are used to characterize combinatorial codes of odorant identities at the OSN level (middle) and the PN level (right)

Detailed connectivity information informed the model construction for both the adult and larva EOS, that we developed based on parameterized versions of the previous literature [14].

In particular, the larval model includes fewer number of OSNs, PNs and LNs in Antenna and Antennal Lobe circuit as shown in Fig. 4(b) right.

The adult and larval EOS models were simultaneously evaluated on a collection of mono-molecular odorants whose binding affinities to odorant receptors have been estimated from physiological recordings (see also Methods). In Figure 4(c) (left), the affinity values are shown for the odorant receptors that are only in the adult fruit fly (top panel), that appear in both the adult and the larva (middle panel) and, finally, that are only in the larva. The steady-state responses of the Antenna and Antennal Lobe circuit for both models are computed and shown in Figure.4 (c) (middle and right, respectively). Visualized in juxtaposition alongside the corresponding affinity vectors, we observe stark contrast in odorant representation at all layers of the circuit between adult and larva, raising the question of how downstream circuits can process differently represented odorant identities and instruct similar olfactory behavior across development. Settling such questions require additional physiological recordings, that may improve the accuracy of the current FlyBrainLab EOS circuit models.

## Discussion

Historically, a large number of visualization and computational tools have been developed primarily designed for either neurobiological studies (see Figure 1 (left) or computational studies (see Figure 1 (right)). These are briefly discussed below.

The computational neuroscience community has invested a significant amount of effort in developing tools for analyzing and evaluating model neural circuits. A number of simulation engines have been developed, including general simulators such as NEURON [17], NEST [18], Brian [19], Nengo [20], Neurokernel [6], DynaSim [21], and the ones that specialize in multi-scale simulation, *e*.*g*., MOOSE [22], in compartmental models, *e*.*g*., ARBOR [23], and in fMRI-scale simulation *e*.*g*., Virtual Brain [24, 25]. Other tools offer to improve the accessibility of these simulators by facilitating the creation of large-scale neural networks, e.g., BMTK [26] and NetPyNe [27], and by providing a common interface, simplifying the simulation workflow and streamlining parallelization of simulation, *e*.*g*., PyNN [28], Arachne [29] and NeuroManager [30]. To facilitate access and exchange of neurobiological data worldwide, a number of model specification standards have been worked upon in parallel including MorphML [31], NeuroML [32], SpineML [33] and SONATA (https://github.com/AllenInstitute/sonata).

Even with the help of these computational tools, it still takes a substantial amount of manual effort to build executable circuits from real data provided, for example, by model databases such as ModelDB/NeuronDB [34] and NeuroArch [5]. Moreover with the ever expanding size of the fly brain datasets, it has become more difficult to meet the demand of creating executable circuits that can be evaluated with different datasets. In addition, with very few exceptions, comparisons of circuit models, a standard process in the computer science community, are rarely available in the computational neuroscience literature.

Substantial efforts by the system neuroscience community went into developing tools for visualizing the anatomy of the brain. A number of tools have been developed to provide interactive, web-based interfaces that allow neurobiologists to extensively explore and visualize neurobiological information of the fruit fly brain and ventral nerve cord, for both the adult [1, 4] and the larva [2, 35]. These include the FlyCircuit [1], the Fruit Fly Brain Observatory (FFBO/NeuroNLP) [7], Virtual Fly Brain [36], neuPrintExplorer [37], and CATMAID [38]. Similar tools have been developed for other model organisms, such as the Allen Mouse Brain Connectivity Atlas [39], the WormAtlas for *C*. *elegans* (http://www.wormatlas.org) and the Z Brain for zebra fish [40]. A number of projects, *e*.*g*., [41], offer a more specialized capability for visualizing and analyzing neuroanatomy data.

Although primarily designed for guidance in system experimentation, these visualization tools have significantly improved the access to and exploration of brain data. A number of these efforts started to bridge the gap between neurobiological data and computational modeling including the Geppetto [42], the OpenWorm [43] and the Open Source Brain [44] initiatives and the Brain Simulation Platform of the Human Brain Project [45].

Without information linking circuit activity/computation to the structure of the underlying neuronal circuits, understanding the function of brain circuits remains, however, elusive.

Lacking a systematic method of automating the process of creating and exploring the function of executable circuits at the brain or system scale levels hinders the application of these tools when composing more and more complex circuits. Furthermore, these tools fall short of offering the capability of generating static circuit diagrams, let alone interactive ones. The experience of VLSI design, analysis and evaluation of computer circuits might be instructive here. An electronic circuit engineer reads a circuit diagram of a chip, rather than the 3D structure of the tape-out, to understand its function, although the latter ultimately realizes it. Similarly, visualization of a biological circuit alone, while powerful and intuitive for building a neural circuit, provides little insights into the function of the circuit. While simulations can be done without a circuit diagram, understanding how an executable circuit leads to its function remains elusive.

The tools discussed above all fall short of offering an integrated infrastructure that can effectively leverage the ever expanding connectomic/synaptomic and neurophysiology data for creating and exploring executable fly brain circuits. Creating circuit simulations from visualized data remains a major challenge and requires extraordinary effort in practice as amply demonstrated by the Allen Brain Observatory [46]. The need to accelerate the pace of discovery of the functional logic of the brain of model organisms has entered a center stage in brain research.

FlyBrainLab is uniquely positioned to accelerate the discovery of the functional logic of the *Drosophila* brain. Its interactive architecture seamlessly integrates and brings together computational models with neuroanatomical, neurogenetic and electrophysiological data, organizing neuroscientific fly brain data from a group of seemingly disparate databases, arrays and tables, into a well structured data and executable circuit repository.

As detailed here, the FlyBrainLab UI supports a highly intuitive and automated workflow that streamlines the 3D exploration and visualization of fly brain circuits, and the interactive exploration of the functional logic of executable circuits created directly from the explored and visualized fly brain data. In conjunction with the capability of visually constructing circuits, speeding up the process of creating interactive executable circuit diagrams can substantially reduce the exploratory development cycle.

The FlyBrainLab Utility and Circuit Libraries accelerate the creation of models of executable circuits. The Utility Libraries (see also Supplementary Section 2) help untangle the graph structure of neural circuits from raw connectome and synaptome data. The Circuit Libraries (see also Supplementary Section 3) facilitate the exploration of neural circuits of the neuropils of the central complex and, the development and implementation of models of the adult and larva fruit fly early olfactory system.

Importantly, to transcend the limitations of the connectome, FlyBrainLab is providing Circuit Libraries for molecular transduction in sensory coding (see also Supplementary Section 3), including models of sensory transduction and neuron activity data. These libraries serve as entry points for discovery of circuit function in the sensory systems of the fruit fly. They also enable the biological validation of developed executable circuits within the same platform.

The modular software architecture underlying FlyBrainLab provides substantial flexibility and scalability for the study of the larva and adult fruit fly brain. As more data becomes available, we envision that the entire central nervous system of the fruit fly can be readily explored with FlyBrainLab. Furthermore, the core of the software and the workflow enabled by the FlyBrainLab for accelerating discovery of *Drosophila* brain functions can be adapted in the near term to other model organisms including the zebrafish and bee.

## Acknowledgments

The research reported here was supported by AFOSR under grant #FA9550-16-1-0410 and DARPA under contract #HR0011-19-9-0035.

## Author Contributions

A.A.L., T.L., M.K.T. and Y.Z. conceived the study and FlyBrainLab software architecture. T.L. and M.K.T. and Y.Z. developed the FlyBrainLab platform. T.L. and M.K.T. developed user-side libraries, Y.Z. updated the server-side components of the existing FFBO architecture, M.K.T. developed utility libraries. A.A.L. and Y.Z. developed comparative models of the central complex. A.A.L., T.L. and M.K.T. developed comparative models of the early olfactory system. A.A.L., T.L., M.K.T and Y.Z. wrote and revised the manuscript.

## Methods

The FlyBrainLab interactive computing platform tightly integrates tools enabling the morphological visualization and exploration of large connectomics/synaptomics datasets, interactive circuit construction and visualization and multi-GPU execution of neural circuit models for *in silico* experimentation. The tight integration is achieved with a comprehensive open software architecture and libraries to aid data analysis, creation of executable circuits and exploration of their functional logic.

### Architecture of FlyBrainLab

FlyBrainLab exhibits a highly extensible, modularized architecture consisting of a number of interconnected server-side and user-side components (see Supplementary Figure **1**) including the NeuroArch Database, the Neurokernel Execution Engine and the NeuroMinerva front-end. The architecture of FlyBrainLab and the associated components are described in detail in the Supplementary Information Section 1.

### FlyBrainLab Utilities Libraries

FlyBrainLab offers a number of utility libraries to untangle the graph structure of neural circuits from raw connectome and synaptome data. These libraries provide a large number of tools including high level connectivity queries and analysis, algorithms for discovery of connectivity patterns, circuit visualization in 2D or 3D and morphometric measurements of neurons. These utility libraries are described in detail in the Supplementary Information Section 2.

### FlyBrainLab Circuit Libraries

FlyBrainLab provides a number of libraries for analysis, evaluation and comparison of fruit fly brain circuits. The initial release of FlyBrainLab offers libraries for exploring neuronal circuits of the central complex, early olfactory system, and implementations of olfactory and visual transduction models. These circuit libraries are described in detail in the Supplementary Information Section 3.

### Analyzing, Evaluating and Comparing Circuit Models of the Fruit Fly Central Complex

Model A [8], Model B [9] and Model C [10] were implemented using the CXcircuit Library (see also Supplementary Information Section 3). The circuit diagram of the wild-type fruit fly provided by the CXcircuit Library underlies model A. It is built by querying all the neurons of the CX in the FlyCircuit dataset. The innervation pattern of each neuron was visually examined in the NeuroNLP window and a standard name assigned according to the naming scheme adopted in the CXcircuit Library. The neurons with missing morphologies in the FlyCircuit dataset were completed by referencing data available in the literature [47, 48]. Construction of the models B and C were obtained by interactively removing and adding neurons in the circuit diagram of model A.

All three models include the same core subcircuits for modeling the Protocerebral Bridge - Ellipsoid Body (PB-EB) interaction. The core subcircuits include 3 cell types, namely, the PB-EB-LAL, PB-EB-NO and EB-LAL-PB neurons (NO - Noduli, LAL - Lateral Accessory Lobe, see also [8] for a list of synonyms of each neuron class). These cells innervate three neuropils, either PB, EB and LAL or PB, EB, and NO. Note that only synapses within PB and EB are considered. For model A, this is achieved by removing all neurons that do not belong to the core PB-EB circuit. Model B includes an additional cell type, the PB local neurons that introduce global inhibition to the PB-EB circuit. Model C does not include PB local neurons, but models 3 types of ring neurons that innervate the EB. Both PB local neurons and ring neurons are present in model A. However, except for their receptive fields, all ring neurons in model A are the same (see below). Figure 5 depicts the correspondence established between the morphology of example neurons and their respective representation in the overall circuit diagram.

**Figure 5:**
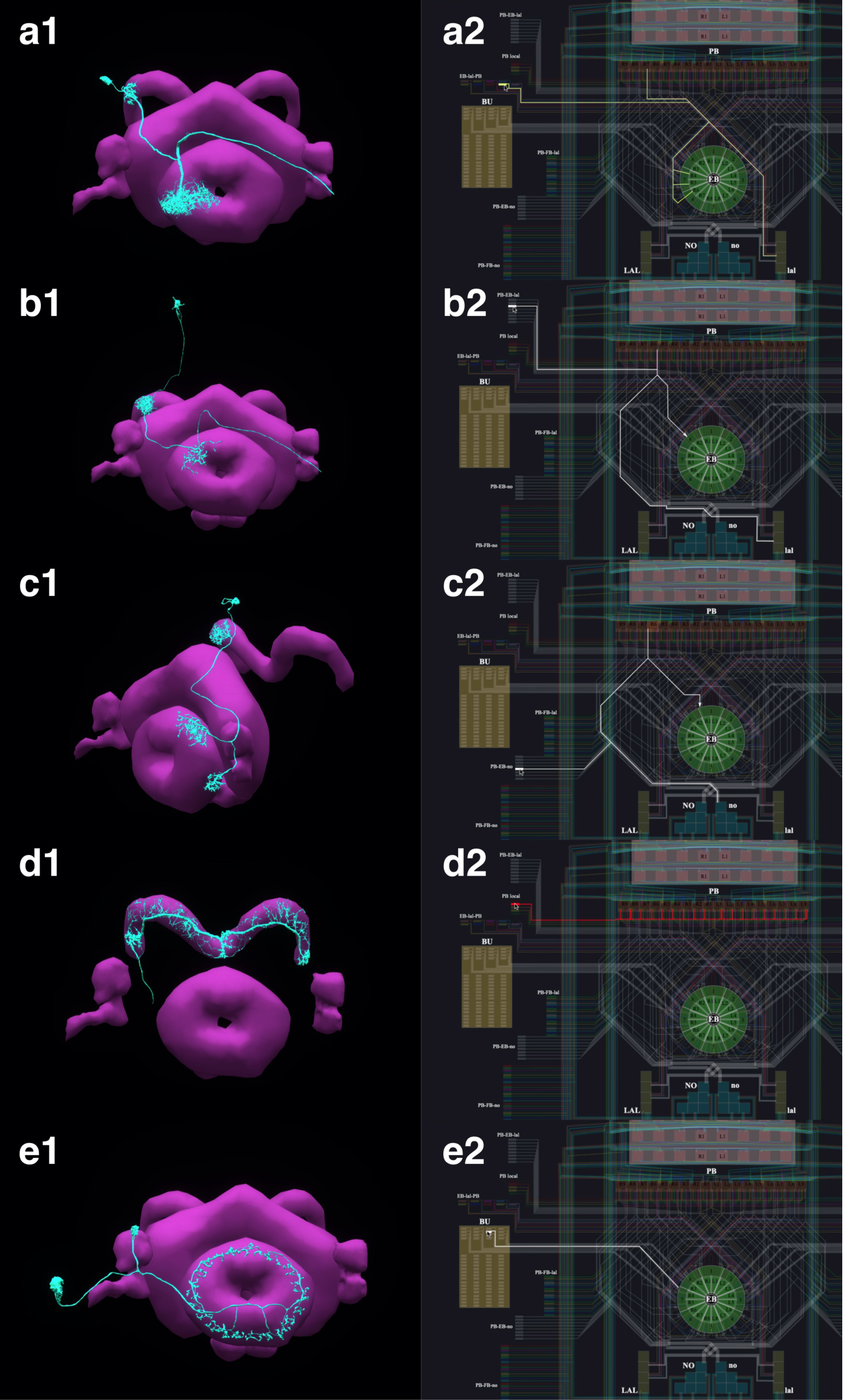
The correspondence between the morphology and the circuit diagram representation of 5 classes of neurons that determine the PB-EB interaction. (a1, a2) EB-LAL-PB neuron and its wiring in the circuit diagram. (b1, b2) PB-EB-LAL neuron and its wiring in the circuit diagram. (c1, c2) PB-EB-NO neuron and its wiring in the circuit diagram. (d1, d2) PB local neuron and its wiring in the circuit diagram. (e1, e2) Ring neuron and its wiring in the circuit diagram.

In Model C, the subcircuit consisting of the PB-EB-LAL and EB-LAL-PB neurons was claimed to give rise to the persistent bump activity while the interconnect between PB-EB-NO and EB-LAL-PB allowed the bump to shift in darkness. To better compare with the other two models that did not model the shift in darkness, PB-EB-NO neurons were interactively disabled from the diagram.

For a single vertical bar presented to the fly at the position shown in Figure 2(d1) (see also the flattened visual input in Figure 2(d2)), the PB glomeruli or the EB wedges innervated by neurons of each of the 3 circuit models that receive injected current or external spike inputs are, respectively, highlighted in Figure 2(a3, b3, c3). The CXcircuit Library generates a set of visual stimuli and computes the spike train and/or the injected current inputs to each of the 3 models.

For model A [8], each PB glomerulus is endowed with a rectangular receptive field that covers 20*°* in azimuth and the entire elevation. Together, the receptive fields of all PB glomeruli tile the 360*°* azimuth. All PB neurons with dendrites in a glomerulus, including the PB-EB-LAL and PB-EB-NO neurons, receive the visual stimulus filtered by the receptive field as an injected current. Additionally, each Bulb (BU) microglomerulus is endowed with a Gaussian receptive field with a standard deviation of 9*°* in both azimuth and elevation. The ring neuron innervating a microglomerulus receives the filtered visual stimulus as an injected current (see also the arrows in Figure 2(a3)). Neuron dynamics follow the Leaky Integrate-and-Fire (LIF) neuron model

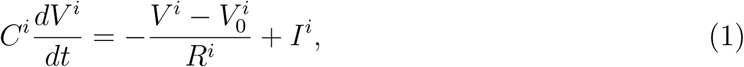

where *V* ^*i*^ is the membrane voltage of the *i*th neuron, *C*^*i*^ is the membrane capacitance, 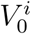 is the resting potential, *R*^*i*^ is the resistance, and *I*^*i*^ is the synaptic current. Upon reaching the threshold voltage 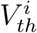, each neuron’s, membrane voltage is reset to 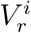. Synapses are modeled as *α*-synapses with dynamics given by the differential equations

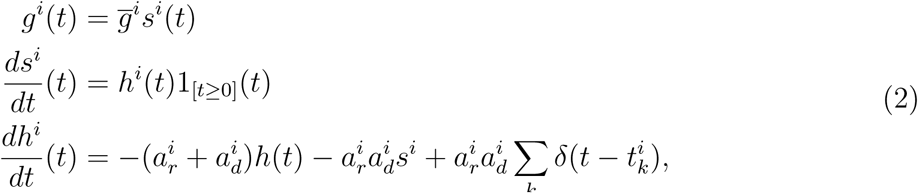

where 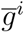 is a scaling factor, 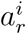 and 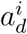 are, respectively, the rise and decay time of the synapse, 1_[*t≥*0]_(*t*) is the Heaviside function and *δ*(*t*) is the Dirac function. 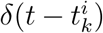 indicates an input spike to the *i*th synapse at time 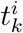.

For Model B [9], the receptive field of each of the 16 EB wedges covers 22.5*°* in azimuth. All EB-LAL-PB neurons that innervate a wedge receive a spike train input whose rate is proportional to the filtered visual stimulus (see also arrow in Figure 2(b3)). The maximum input spike rate is 120 Hz when the visual stimulus is a bar of width 20*°* at maximum brightness 1. A 5 Hz background firing is always added even in darkness. Neurons are modeled as LIF with a refractory period of 2.2 ms as suggested in [9]. For synapses, instead of using the postsynaptic current (PSC)-based model described in [9], we used the *α*-synapse described above and chose the parameters such that the time-to-peak and peak value approximately matched that of the PSC-based synapse.

For Model C [10], the receptive field of each of the 16 EB wedges covers 22.5*°* in azimuth. Two PB-EB-LAL neurons project to a wedge each from a different PB glomerulus. Input to the Model C circuit is presented to pairs of PB glomeruli (see also arrows in Figure 2(c3)), and all neurons with dendrites in these two PB glomeruli receive a spike train input at a rate proportional to the filtered visual stimuli, with a maximum 50 Hz when the bar is at maximum brightness 1. Neurons are modeled as LIF neurons with a refractory period of 2 ms (as suggested in [10]). Synapses are either modeled by the AMPA/GABA_*A*_ receptor dynamics as

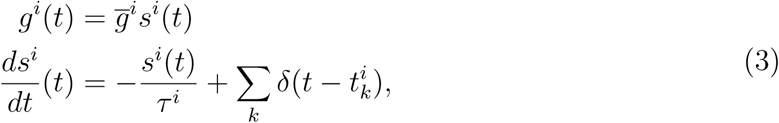

where *g*^*i*^(*t*) is the synaptic conductance, *τ*^*i*^ is the time constant, and *s*^*i*^(*t*) is the state variable of the *i*th synapse, or modeled by the NMDA receptor dynamics [10]

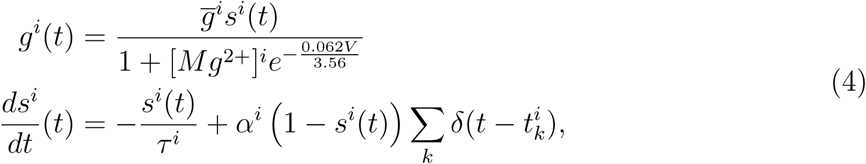

where *g*^*i*^(*t*) is the synaptic conductance, 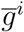 is the maximum conductance, *s*^*i*^(*t*) is the state variable, *τ*^*i*^ is the time constant, *α*^*i*^ *>* 0 is a constant, [*Mg*^2+^]^*i*^ is the extracellular concentration of *Mg*^2+^, respectively, of the *i*th synapse and *V* is the membrane voltage of the postsynaptic neuron.

Parameters of the above models can be found in [8–10] and in the CXcircuit Library.

The 35-second visual stimulus, depicted in Figure 2(d3), was presented to all 3 models. A bright vertical bar moves back and forth across the entire visual field while a second bar with lower brightness is presented at a fixed position. Figure 2(d3, bottom) depicts the time evolution of the visual input.

To visualize the response of the 3 executable circuits, the mean firing rate *r*^*j*^(*t*) of all EB-LAL-PB neurons that innervate the *j*th EB wedge was calculated following [10]

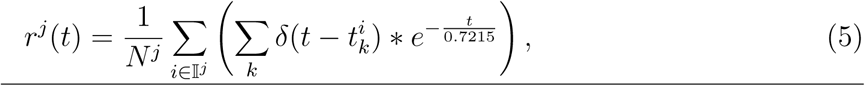

where *∗* denotes the convolution operator, 𝕀^*j*^ is the index set of EB-LAL-PB neurons that innervate the *j*th EB wedge, whose cardinality is *N*^*j*^, and 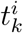 is the time of *k*th spike generated by *i*th neuron. CXcircuit Library provides utilities to visualize the circuit response.

### Analyzing, Evaluating and Comparing of Adult Antenna and Antennal Lobe Circuit Models Based upon the FlyCircuit and Hemibrain Datasets

The Early Olfactory System models based on the FlyCircuit and the Hemibrain datasets were implemented using the EOScircuits Library (see also Supplementary Information Section 3). The circuit architecture, shown in Figure 3(b, left), follows previous work [49] based upon the anatomically observed connectivity between local neurons (LN) and projection neurons (PNs) (see also Figure 3(a, left)). At the input layer (the Antenna Circuit), the stimulus model for the adult EOS circuit builds upon affinity data from the DoOR dataset [50], with physiological recordings for 23/51 receptor types. Receptors for which there was no affinity data in the DoOR dataset were assumed to have zero affinity values. Two input stimuli were used. The first input stimulus is 5-second long, and between 1-3 second, ammonium hydroxide with a constant concentration of 100 ppm was presented to the circuits in Figure 3(b) and the responses are shown in (Figure 3(c)). The same odorant waveform is used here as in Figure 3(d). To generate the concentration waveform of the odorant, values were drawn randomly from the uniform distribution between 0 and 100 ppm every 10^*−*4^ seconds between 1-3 seconds in Figure 3(d). The sequence is then filtered by a lowpass filter with a 30Hz bandwidth [15] to obtain the concentration of the odorant.

Olfactory Sensory Neurons expressing each one receptor type processed the input odorant in parallel. The Antennal Lobe model based on FlyCircuit data is divided into two sub-circuits: 1) the ON-OFF circuit and 2) the Predictive Coding circuit [16]. The ON-OFF circuit describes odorant gradient encoding by Post-synaptic LNs in the AL, while the Predictive Coding circuit describes a divisive normalization mechanism by Pre-synaptic LNs that enable concentration-invariant odorant identity encoding by Projection Neurons in the AL.

The EOS model based on Hemibrain dataset takes advantage of the detailed connectivity between neurons (see Figure 3(a, right) and introduces a more extensive connectome-synaptome model of the AL (see Figure 3(b, right)). FlyBrainLab utility libraries were used to 1) access the Hemibrain data, 2) find PNs and group them by glomeruli, 3) use this data to find the OSNs associated with each glomerulus, 4) find LNs and group connectivity between OSNs, LNs and PNs. We constructed in FlyBrainLab an executable circuit using the Hemibrain data. In addition to the baseline model in Figure 3(b, left), we introduced 1) LNs that innervate specific subsets of glomeruli, 2) LNs that provide inputs to both OSN axon terminals and to PNs dendrites, 3) synapses from PNs onto LNs.

### Evaluating, Analyzing and Comparing Early Olfactory Circuit Models of the Larva and the Adult Fruit Flies

The Early Olfactory System models for both the adult and the larval flies were implemented using the EOScircuits library (see also Supplementary Information Section 3). The circuit of the adult EOS follows the one described above. Similarly, the larval model is implemented using physiological recording on 14/21 receptor types [51]. In both the adult and larval physiology datasets, 13 common mono-molecular odorants were employed (see Figure 4(d legend). Together, 13/23 odorant/receptor pairs for adult and 13/14 odorant/receptor pairs for larva were used for model evaluation, where each odorant was carried by a 100 ppm concentration waveform. In both adult and larva Antenna circuits, Olfactory Sensory Neurons expressing each receptor type processed an odorant waveform in parallel.

The adult Antennal Lobe model follows the one built on the Hemibrain data [4]. Both the adult and the larva circuit components are parameterized by the number of LNs per type, where for instance there were 28 LNs used in the larval model in accordance to connectome data [2]. In addition to neuron types, the AL circuit was modified in terms of connectivity from 1) LNs to Projection Neurons (PNs), 2) PNs to LNs and 3) LNs to LNs. The evaluation of both EOS models focused on the Input/Output relationship comparison between the adult and the larval EOS models. For each of the 13 odorants, the input stimulus is a 5 second concentration waveform that is 100ppm from 1-4 second and 0 ppm otherwise. Both adult and larval models reach steady-state after 500ms and the steady-state population responses averaged across 3-4 seconds are computed as odorant combinatorial code at each layer (i.e., OSN response, PN response).

### Code Availability and Installation

Stable and tested FlyBrainLab installation instructions for user-side components and utility libraries are available at https://github.com/FlyBrainLab/FlyBrainLab for Linux, MacOS and Windows. The installation and use of FlyBrainLab does not require a GPU, but a service-side backend must be running, for example, on a cloud service, that the user-side of FlyBrainLab can connect to. The serverside backend codebase is publicly available at https://github.com/fruiflybrain and https://github.com/neurokernel.

A full installation of FlyBrainLab, including all backend and frontend components, is available as a Docker image at https://hub.docker.com/r/fruitflybrain/fbl. The image requires a Linux host with at least 1 CUDA-enabled GPU and the nvidia-docker package (https://github.com/NVIDIA/nvidia-docker) installed. For a custom installation of the complete FlyBrainLab platform, a shell script is available at https://github.com/FlyBrainLab/FlyBrainLab.

### Data Availability

The NeuroArch Database created from publicly available FlyCircuit, Hemibrain and Larva L1EM datasets can be downloaded from https://hub.docker.com/r/fruitflybrain/fbl.

## Supplementary Information

### 1 The Architecture of FlyBrainLab

To support the study of the function of brain circuits FlyBrainLab implements an extensible, modularized architecture that tightly integrates fruit fly brain data and models of executable circuits. Supplementary Figure **1** depicts the architecture of FlyBrainLab on both the userand backend server-side.

The backend server-side components are described in section 1.1. The user interface and user-side components are presented in section 1.2.

#### 1.1 The Server-Side Components

The server-side backend consists of 4 components: FFBO Processor, NeuroArch, Neurokernel and NeuroNLP servers. They are collectively called the FFBO servers. A brief description of each of the components is given below.

##### FFBO Processor

implements a Crossbar.io router (https://crossbar.io/) that establishes the communication path among connected components. Components communicate using routed Remote Procedure Calls (RPCs) and a publish/subscribe mechanism. The routed RPCs enable functions implemented on the server-side to be called by the user-side backend (see also Section 1.2). After an event occurs, the publisher immediately informs topic subscribers by invoking the publish/subscribe mechanism. This enables, for example, the FFBO processor to inform the user-side and other servers when a new backend server is connected. The FFBO processor can be hosted locally or in the cloud. It can also be hosted by a service provider for, *e*.*g*., extra data sharing. The open source code of the FFBO processors is available at https://github.com/fruitflybrain/ffbo.processor.

##### NeuroArch Server

hosts the NeuroArch graph database [1] implemented with OrientDB (https://orientdb.com). The NeuroArch Database provides a novel data model for representation and storage of connectomic, synaptomic, cell type, activity and genetic data of the fruit fly brain with cross-referenced executable circuits. The NeuroArch data model is the foundation of the integration of fruit fly brain data and executable circuits in FlyBrainLab. Low-level queries of the NeuroArch Database are supported by the NeuroArch Python API (https://github.com/fruitflybrain/neuroarch). The NeuroArch Server provides high level RPC APIs for remote access of the NeuroArch Database. The open source code of the NeuroArch Server is available at https://github.com/fruitflybrain/ffbo.neuroarch_component.

##### Neurokernel Server

provides RPC APIs for code execution of model circuits by the Neurokernel Execution Engine [2]. Neurokernel supports the easy combination of independently developed executable circuits towards the realization of a complete whole brain emulation. The Neurokernel Execution Engine features:

- the core Neurokernel services (https://github.com/neurokernel/neurokernel) providing management capabilities for code execution, and communication between interconnected circuits,
- the Neurodriver services (https://github.com/neurokernel/neurodriver) providing low level APIs for code execution on GPUs according to user-specified circuit connectivity, biological spike generators and synapses, and input stimuli.

The Neurokernel Server directly fetches the specification of executable circuits from the NeuroArch Server, instantiates these circuits and transfers them for execution to the Neurokernel Execution Engine. The open source code of the Neurokernel Server is available at https://github.com/fruitflybrain/ffbo.neurokernel_component.

##### NeuroNLP Server

provides an RPC API for translating queries written as English sentences, such as “add dopaminergic neurons innervating the mushroom body”, into database queries that can be interpreted by the NeuroArch Server API. This capability increases the accessibility of the NeuroArch Database to researchers without prior exposure to database programming, and facilitates research by simplifying the often-demanding task of writing database queries. The open source code of the NeuroNLP Server is available at https://github.com/fruitflybrain/ffbo.nlp_component.

**Supplementary Figure 1:**
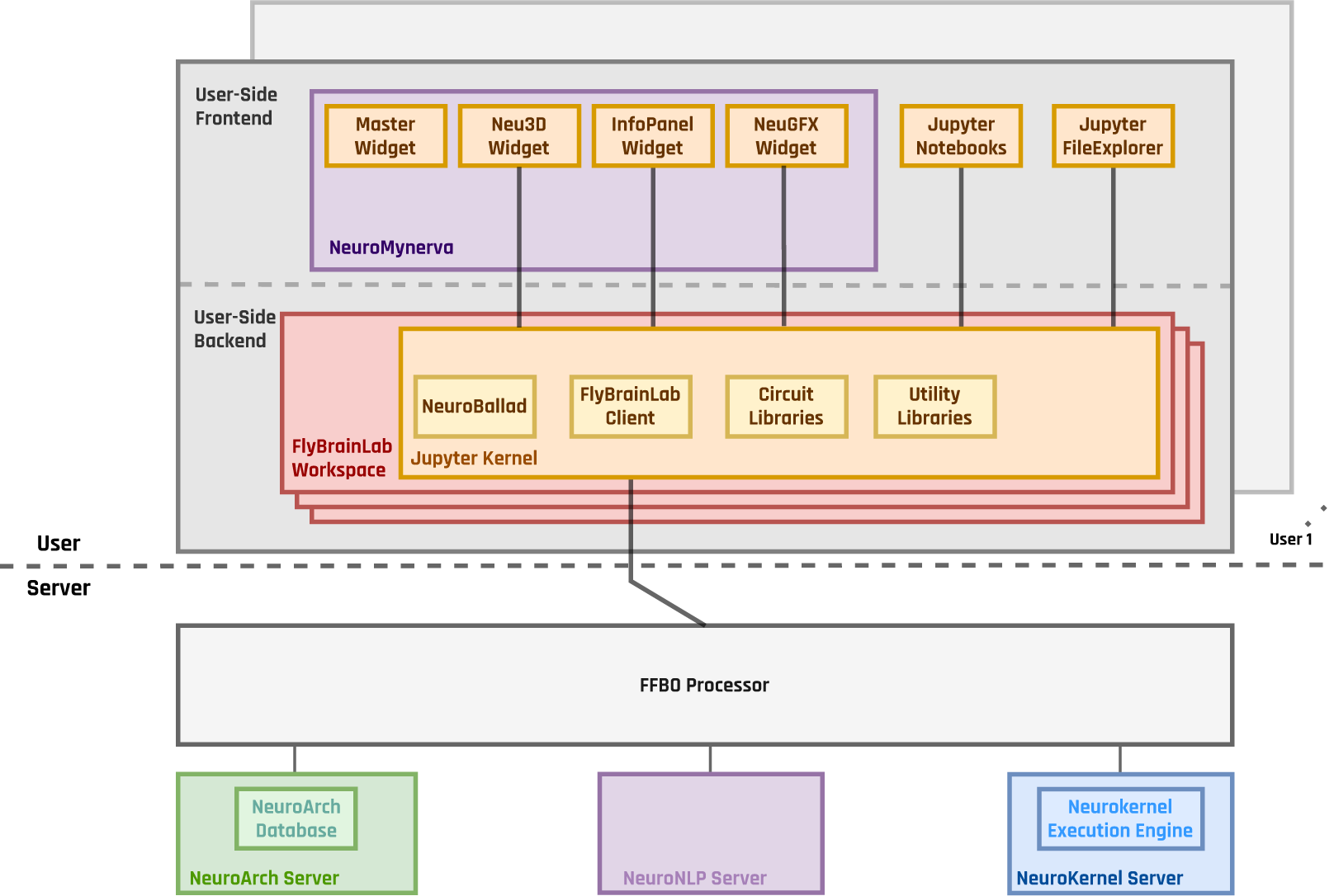
The Architecture of FlyBrainLab. The server-side architecture [3] consists of the FFBO Processor, the NeuroNLP Server, the NeuroArch Server and the Neurokernel Server. The user-side provides the local execution environment as well as an easy-to-use GUI for multi-user access to the services provided by the server-side. The FlyBrainLab Utility Libraries and Circuit Libraries (see Section 2 and 3 for details) can be loaded into the FlyBrainLab workspace of the user-side backend.

#### 1.2 The User Interface and User-Side Components

The FlyBrainLab user-side consists of the NeuroMynerva frontend and the FlyBrainLab Client and Neuroballad backend components. A brief description of each of the components is given below.

##### NeuroMynerva

is the user-side frontend of FlyBrainLab. It is a browser-based application that substantially extends upon JupyterLab by providing a number of widgets, including a Neu3D widget for 3D visualization of fly brain data, a NeuGFX widget for exploring executable neural circuits with interactive circuit diagrams, and an InfoPanel widget for accessing individual neuron/synapse data. All widgets communicate with and retrieve data from the FlyBrainLab Client. A master widget keeps track of the instantiated widgets by the user interface. With the JupyterLab native notebook support, APIs of the FlyBrainLab Client can be directly called from notebooks. Such calls provide Python access to NeuroArch queries and circuit execution. Interaction between code running in notebooks and widgets is fully supported.

##### FlyBrainLab Client

is a user-side backend implemented in Python that connects to the FFBO processor and accesses services provided by the connected backend servers. FlyBrainLab Client provides program execution APIs for handling requests to the server-side components and parsing of data coming from backend servers. The FlyBrainLab Client also exhibits a number of high-level APIs for processing data collected from the backend servers, such as computing the adjacency matrix from retrieved connectivity data or retrieving morphometrics data. In addition, it handles the communication with the frontend through the Jupyter kernel.

##### NeuroBallad

is a Python library that simplifies and accelerates executable circuit construction and simulation using Neurokernel in Jupyter notebooks in FlyBrainLab. NeuroBallad provides classes for specification of neuron or synapse models with a single line of code and contains functions for adding and connecting these circuit components with one another. NeuroBallad also provides capabilities for compactly specifying inputs to a circuit on a per-experiment basis.

##### NeuroMynerva User Interface

(UI) is depicted in the Supplementary Figure **2**. The UI typically consists 5 blocks, including (1) a NeuroNLP 3D visualization window with a search bar for NLP queries, displaying and interacting with fly brain data such as the morphology of neurons and position of synapses, that are all managed by the Neu3D widget; (2) a NeuroGFX executable circuits window, for exploring executable neural circuits with interactive circuit diagrams, that are managed by the NeuGFX widget; (3) a Program Execution window with a built-in Jupyter notebook, executing any Python code including calls to the FlyBrainLab Client, for direct access to database queries, visualization, and circuit execution, (4) an Info Panel displaying details of highlighted neurons including the origin of data, genetic information, morphometric statistics and synaptic partners, etc, all managed by the InfoPanel widget; (5) a Local File Access Panel with a built-in Jupyter file browser for accessing local files.

**Supplementary Figure 2:**
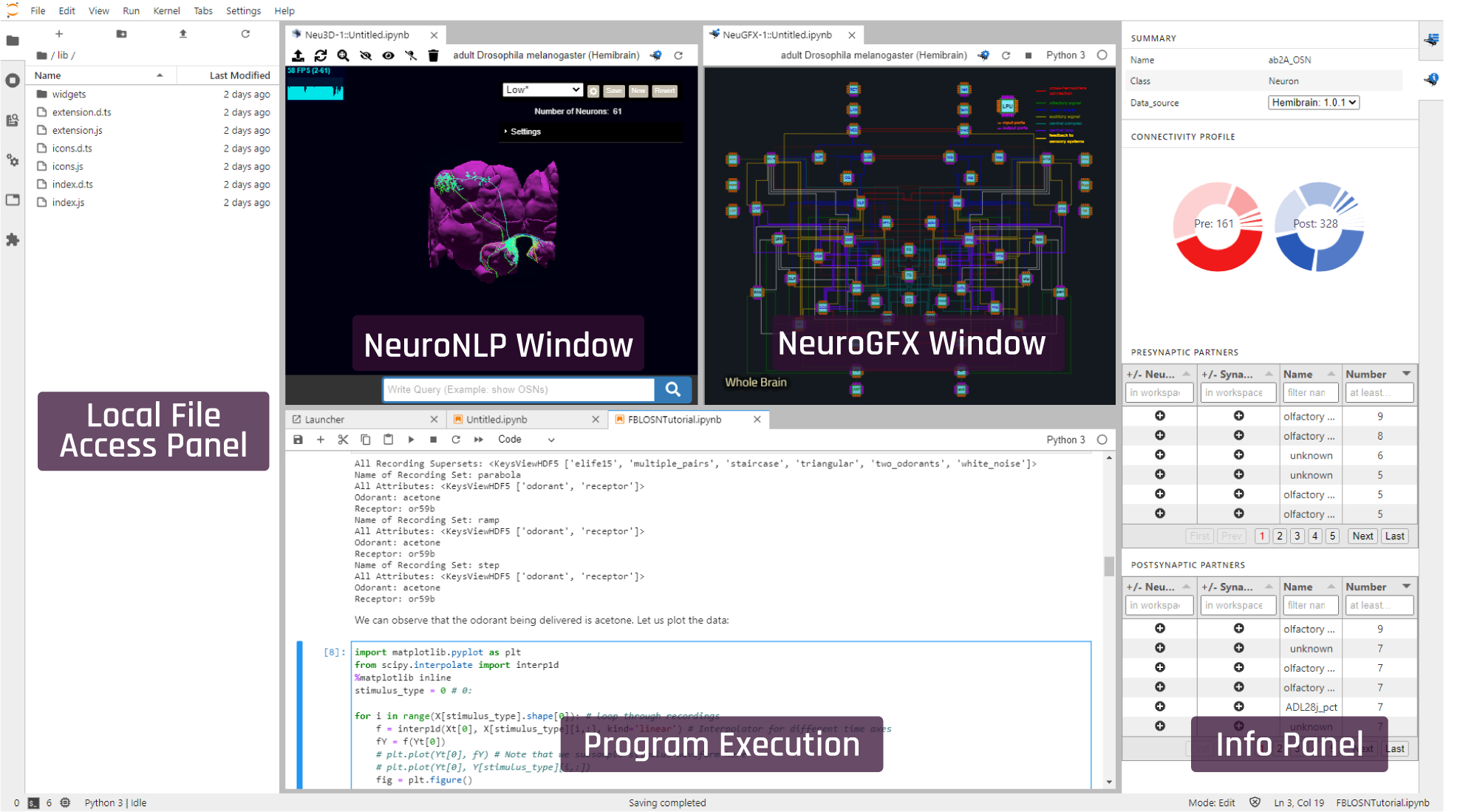
The NeuroMynerva User Interface. (top left) NeuroNLP 3D visualization window. (top right) NeuroGFX executable circuits window. (bottom center) Program Execution window with Jupyter notebook. (right) Information Panel for individual neurons/synapses. (left) Local File Access Panel with Jupyter FileBrowser widget.

### 2 Utility Libraries for the Fruit Fly Connectome/ Synaptome

Different connectome and synaptome datasets are often available at different levels of abstraction. For example, some datasets come with cell types labeled and some only provide raw graph level connectivity. Without extensive analysis tools, it takes substantial manual effort to construct and test a neural circuit. FlyBrainLab offers a number of utilities to explicate the graph structure of neural circuits from raw connectome and synaptome data. In conjunction with the capability of visually constructing circuits enabled by the NeuroMynerva front-end, speeding up the process of creating interactive executable circuit diagrams can substantially reduce the exploratory development cycle.

The FlyBrainLab Utility Libraries include:

- **NeuroEmbed**: Cell Classification and Cell Type Discovery,
- **NeuroSynapsis**: High Level Queries and Analysis of Connectomic and Synaptomic Data,
- **NeuroGraph**: Connectivity Pattern Discovery and Circuit Visualization Algorithms,
- **NeuroWatch**: 3D Fruit Fly Data Visualization in Jupyter Notebooks,
- **NeuroMetry**: Morphometric Measurements of Neurons.

In this section, we outline the capabilities enabled by the Utility Libraries listed above.

#### NeuroEmbed: Cell Classification and Cell Type Discovery

The NeuroEmbed library implements a set of algorithms for structure discovery based on graph embeddings into low-dimensional spaces providing capabilities for:

- Cell type classification based on connectivity, and optionally morphometric features,
- Searching for neurons that display a similar connectivity pattern,
- Standard evaluation functions for comparison of embedding algorithms on clustering and classification tasks.

#### NeuroSynapsis: High Level Queries and Analysis of Connectomic and Synaptomic Data

The NeuroSynapsis Library offers a large set of utilities to accelerate the construction of circuits and analysis of connectomic and synaptomic data. It provides capabilities for

- Retrieval of neuron groups according to user-defined criteria, such as cell type, innervation pattern and connectivity, etc.,
- Retrieval of connectivity between neurons, cell types, or user-defined neuron groups, thought direct or indirect connections,
- Retrieval of synapse positions and partners for groups of neurons and the capability to filter synapses by brain region, partnership or spatial location,
- Statistical analysis of retrieved synapses, such as the synaptic density in a brain region,

#### NeuroGraph: Connectivity Pattern Discovery and Circuit Visualization Algorithms

The NeuroGraph Library offers a set of tools to discover and analyze any connectivity pattern among cell groups within a circuit. Capabilities include:

- Discovery of connectivity patterns between cell populations by automatic generation of connectivity dendrograms with different linkage criteria (such as Ward or average) [4, 5],
- Analysis of the structure of circuits by community detection algorithms such as Louvain, Leiden, Label Propogation, Girvan-Newman and Infomap,
- Analysis of neural circuit controllability, for example discovery of driver nodes [6],
- Comparing observed connectivity between groups of cells with models of random connectivity.

In addition, the NeuroGraph Library provides utilities to visualize the connectivity of a neural circuit to aid the creation of interactive circuit diagrams. Further capabilities include

- Force-directed layout for the architecture-level graph of the whole brain or circuit-level graph of circuits specified by NeuroSynapsis,
- Semi-automated generation of 2D circuit diagrams from specified connectome datasets either at single-neuron or cell-type scale by separating circuit components into input, local and output populations for layouting.

#### NeuroWatch: 3D Fruit Fly Data Visualization in Jupyter Notebooks

The NeuroWatch Library offers utilities to enable visualization of neuron morphology data using Neu3D in Jupyter Notebook cells. Capabilities include:

- Loading brain regions in 3D mesh format,
- Recoloring, rescaling, repositioning and rotating neuropils, neurons and synapses for visualization,
- Interactive alignment of new neuromorphology datasets into FlyBrainLab widgets.

#### NeuroMetry: Morphometric Measurements of Neurons

Morphometric measurements of neurons can be extracted from neuron skeleton data available in connectome datasets in .swc format. NeuroMetry provides utilities for

- Calculating morphometric measurements of neurons that are compatible with NeuroMopho.org [7], such as total length, total surface area, total volume, maximum euclidean distance between two points, width, height, depth, average diameter, number of bifurcations and the maximum path distance,
- Accessing precomputed measurements in currently available datasets in FlyBrainLab, including FlyCircuit and Hemibrain data.

An application of the morphometric measurements is the estimation of energy consumption arising from spike generation in axon-hillocks [8].

#### Examples

In Supplementary Figure **3**, we briefly present a few examples that illustrate some of the capabilities of the FlyBrainLab Utility Libraries.

In Supplementary Figure **3**(a), we used the NeuroGraph library to analyze the connectivity of neurons in the Hemibrain dataset to identify groups of neurons that internally form dense connections. Here, the Louvain algorithm extracted 8 densely connected neuron groups that correspond to known neuropils and brain regions including the antennal lobe (AL), the mushroom body (MB), the lateral horn (LH), the central complex (CX), the anterior ventrolateral protocerebrum (AVLP) and the superior protocerebrum (SP). The other two correspond to the anterior optic tubercle combined with neurons in the posterior brain (AOTUP) and neurons in the anterior central brain (ACB). A diagram of the connections between these groups of neurons is depicted in Supplementary Figure **3**(b).

In Supplementary Figure **3**(c), neurons innervating the VA1v glomerulus of the antennal lobe (from the VA1v connectome dataset [10]) is visualized with NeuroWatch. To untangle the graph structure of this circuit, we applied the Adjacency Spectral Embedding algorithm [11] of the NeuroGraph library (see Supplementary Figure **3**(d)). The results obtained show that the algorithm is able to correctly separate ipsilateral OSNs, contralateral OSNs, local neurons and projection neurons without an explicit knowledge of the existence of these cell types (clusters marked manually). Looking into outliers that lie far away from their clusters may guide future research inquiries into cell types.

In Supplementary Figure **3**(f), we generated a circuit diagram using NeuroGraph and depicted a lateral horn subcircuit downstream of the V glomerulus projection neurons as well as the neuropils that the lateral horn output neurons (LHONs) project to [12]. The corresponding neuronal circuit is depicted in Supplementary Figure **3**(e).

**Supplementary Figure 3:**
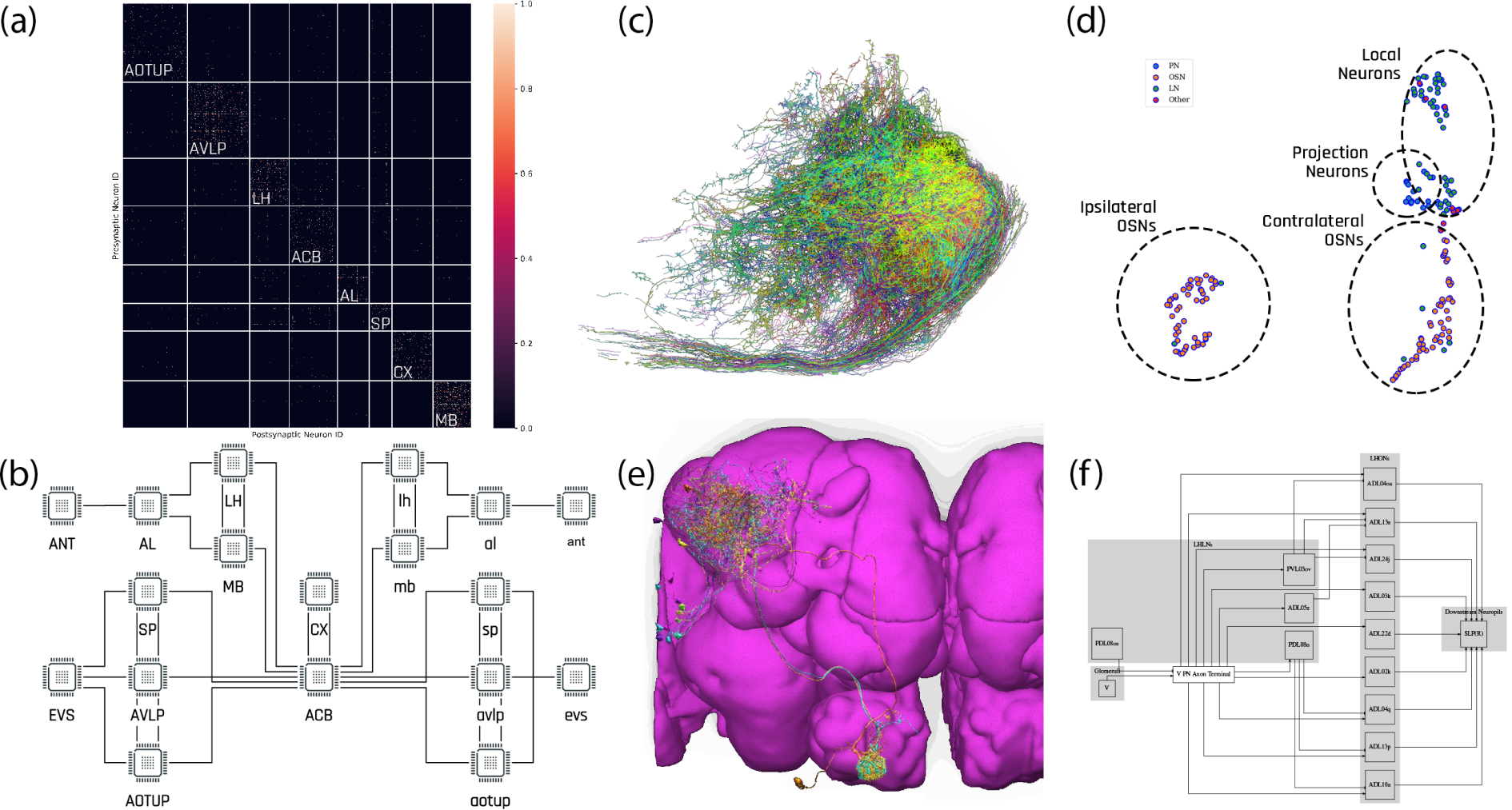
Examples of circuit-level analyses enabled by the FlyBrain-Lab Utility Libraries. **(a)** Louvain algorithm applied to the Hemibrain dataset [9] showing eight groups of densely connected neurons using NeuroGraph. **(b)** A circuit diagram of the *Drosophila* brain drawn using the inter-group edge information for the data in (a). Visual and olfactory inputs from, respectively, the early visual system (EVS) and antenna (ANT) are added. Groups in the left hemisphere added by symmetry. **(c)** VA1v connectomics dataset shown in a Neurowatch window [10]. **(d)** Adjacency Spectral Embedding applied to the VA1v connectomics dataset using the NeuroGraph library shows segregation between local neurons and projection neurons, and suggests connectivity motifs for PNs. **(e)** Uniglomerular projection neurons with dendrites in the V glomerulus and downstream lateral horn neurons visualized with NeuroNLP. **(f)** Circuit diagram automatically generated by the circuit visualization utilities of NeuroGraph starting with the circuit in (e). The diagram shows the circuit downstream of the V glomerulus projection neuron in the lateral horn, and further down to the neuropils that the lateral horn output neurons (LHONs) project to.

### 3 Libraries for Analyzing, Evaluating and Comparing Fruit Fly Brain Circuits

The FlyBrainLab Circuit Library includes:

- **CXcircuits**: Library for Central Complex Circuits,
- **EOScircuits**: Library for Larva and Adult Early Olfactory Circuits,
- **MolTrans**: Library for Molecular Transduction in Sensory Encoding.

These are respectively described in sections 3.1, 3.2 and 3.3 below.

#### 3.1 CXcircuits: Library for Central Complex Circuits

The CXcircuits library facilitates the exploration of neural circuits of the central complex (CX) based on the FlyCircuit dataset [13]. It supports the evaluation and direct comparison of the state-of-the-art of executable circuit models of the CX available in the literature, and accelerates the development of new executable CX circuit models that can be evaluated and scrutinized by the research community at large in terms of modeling assumptions and biological validity. It can be easily expanded to account for the Hemibrain dataset [9].

The main capabilities of the CX Library include programs for:

- constructing *biological* CX circuits featuring
  — a naming scheme for CX neurons that is machine parsable and facilitates the extraction of innervation patterns [14],
  — algorithms for querying CX neurons in the NeuroArch database, by neuron type, by subregions they innervate, and by connectivity,
  — an inference algorithm for identifying synaptic connections between CX neurons in NeuroArch according to their innervation patterns,
- constructing *executable* CX circuit diagrams that
  — are interactive for CX circuits in wild-type fruit flies,
  — interactively visualize neuron innervation patterns and circuit connectivity,
  — are interoperable with 3D visualizations of the morphology of CX neurons,
  — easily reconfigure CX circuits by enabling/disabling neurons/synapses, by enabling/disabling subregions in any of the CX neuropils, and by adding neurons,
  — readily load neuron/synapse models and parameters,
- evaluation of the executable CX circuits with
  — a common set of input stimuli, and the
  — visualization of the execution results with a set of plotting utilities for generating raster plots and a set of animation utilities for creating videos.

### 3.2 EOScircuits Library for Larva and Adult Early Olfactory Circuits

The EOScircuits Library accelerates the development of models of the fruit fly early olfactory system (EOS), and facilitates structural and functional comparisons of olfactory circuits across developmental stages from larva to the adult fruit fly. Built upon FlyBrainLab’s robust execution backend, the EOScircuits Library enables rapid iterative model development and comparison for Antenna (ANT), Antennal Lobe (AL) and Mushroom Body (MB) circuits across developmental stages.

#### ANTcircuits

Modeled after the first layer of the olfactory pathway, the ANTcircuits Library builds upon the Olfactory Transduction (OlfTrans) library (see Section 3.3 below) and describes interactions between odorant molecules and Olfactory Sensory Neurons (OSNs).

The library provides parameterized ANT circuits, that support manipulations including

- Changing the affinity values of each of the odorant-receptor pairs characterizing the input of the Odorant Transduction Process [15],
- Changing parameter values of the Biological Spike Generators (BSGs) associated with each OSN [15],
- Changing the number of OSNs expressing the same Odorant Receptor (OR) type.

#### Alcircuits

Modeled after the second layer of the olfactory pathway, the ALcircuits Library describes the interaction between OSNs in ANT, Projection Neurons in AL and Local Neurons in AL.

The library provides parameterized AL circuits, that support manipulations including

- Changing parameter values of Biological Spike Generators (BSGs) associated with each of the Local and Projection Neurons,
- Changing the number and connectivity of Projection Neurons innervating a given AL Glomerulus,
- Changing the number and connectivity of Local Neurons in the Predictive Coding and ON-OFF circuits of the AL [16].

#### Mbcircuits

Modeled after the third neuropil of the olfactory pathway, the MBcircuits Library describes the expansion-and-sparsification circuit consisting of a population of Antennal Lobe Projection Neurons and Mushroom Body Kenyon Cells (KCs) [17].

The library provides a parameterized the MB subcircuit involving Kenyon Cells and the Anterior Posterior Lateral (APL) neuron, and supports circuit manipulations including

- Generating and changing random connectivity patterns between PNs and KCs with varying degree of fan-in ratio (number of PNs connected to a given KC),
- Changing the strength of feedback inhibition of the APL neuron.

### 3.3 MolTrans Library for Molecular Transduction in Sensory Encoding

The Molecular Transduction Library accelerates the development of models of early sensory systems of the fruit fly brain by providing 1) implementations of transduction on the molecular level that accurately capture the encoding of inputs at the sensory periphery, and 2) activity data of the sensory neurons such as electrophysiology recordings for the validation of executable transduction models.

**The MolTrans Library** includes the following packages:

- **Olfactory Transduction (OlfTrans)**: Molecular Transduction in Olfactory Sensory Neurons,
- **Visual Transduction (VisTrans):** Molecular Transduction in Photoreceptors. The capabilities of the two libraries are discussed in what follows.

#### OlfTrans: Molecular Transduction in Olfactory Sensory Neurons

The OlfTrans Library exhibits the following features and/or capabilities (see also [15]):

- a model of odorant space for olfactory encoding in the adult and larva olfactory system,
- hosts a large number of electrophysiology data of OSNs responding to different odorants with precisely controlled odorant waveforms [18].

Moreover, the OlfTrans Library offers

- the model of an odorant transduction process (OTP) validated by electrophysiology data and executable on Neurokernel/NeuroDriver,
- algorithms for fitting and validation of OTP models with electrophysiology data,
- algorithms for importing odorant transduction models and data into NeuroArch and execution on Neurokernel.

The OlfTrans Library provides critical resources in the study of any subsequent stages of the olfactory system. It serves as an entry point for discovering the function of the circuits in the olfactory system of the fruit fly.

#### VisTrans: Molecular Transduction in Photoreceptors

The VisTrans Library exhibits the following features and/or capabilities (see also [19]):

- a geometrical mapping algorithm of the visual field onto photoreceptors of the retina of the fruit fly,
- a molecular model of the phototransduction process described and biologically validated in [20],
- a parallel processing algorithm emulating the visual field by the entire fruit fly retina,
- algorithms for importing phototransduction models into the NeuroArch Database and for program execution on the Neurokernel Execution Engine.
- algorithms for visually evaluating photoreceptor models.

The VisTrans Library accelerates the study of the contribution of photoreceptors towards the overall spatiotemporal processing of visual scenes. It also serves as an entry point for discovering circuit function in the visual system of the fruit fly.

